# Enhancing daily light exposure increases the antibody response to influenza vaccination in patients with dementia

**DOI:** 10.1101/2020.11.30.405175

**Authors:** Mirjam Münch, Rolf Goldbach, Naomi Zumstein, Petra Vonmoos, Jean-Louis Scartezzini, Anna Wirz-Justice, Christian Cajochen

**Affiliations:** Solar Energy and Building Physics Laboratory, Environmental and Civil Engineering Institute, Ecole Polytechnique Fédérale de Lausanne (Switzerland); Centre for Chronobiology, University Psychiatric Hospitals Basel, Switzerland; Geriatric Service of the City of Zurich, Zurich (Switzerland); Department of Anthropology, McGill University, Montreal (Canada); University Research Priority Program “Dynamics of Healthy Aging”, University of Zurich, Zurich (Switzerland); Sonnweid – The Home, Wetzikon (Switzerland); Transfaculty Research Platform, Molecular and Cognitive Neurosciences, University of Basel (Switzerland)

## Abstract

Enhancing lighting conditions in institutions for individuals with dementia improves their sleep, circadian rhythms and well-being. Here, we tested whether a greater long-term daily light exposure supports the immune response to the annual influenza vaccination. Eighty older institutionalised patients suffering from dementia (54 women and 26 men) continuously wore an activity tracker for 8 weeks to assess individual light exposure and rest-activity cycles. The patients’ immune response was analysed from two blood samples taken before and 4 - 5 weeks after the annual influenza vaccination. Individual antibody concentrations to three influenza virus strains (H3N2, H1N1, IB) were quantified via hemagglutination inhibition assays. By quantifying individual light exposure profiles (including daylight), we classified the patients into a low and a high light exposure group based on a median illuminance of 392.6 lux. The two light exposure groups did not differ in cognitive impairment severity, age or gender distribution. However, patients in the high light exposure group showed a significantly greater circadian rest-activity amplitude (i.e., more daytime activity and less nighttime activity) along with a significantly greater antibody titer increase to the H3N2 vaccine than patients in the low light exposure group, despite similar pre-vaccination concentrations. Sufficient seroprotective responses to all three influenza virus strains were attained for ≥ 75% of participants. These data provide first evidence for a potentially enhanced immune response in patients with dementia when they received more daily light. Increasing daily light exposure may have beneficial effects on the human immune system, either directly or via circadian rhythm stabilisation.

## Introduction

In older patients with dementia, the decline of cognitive functions is often accompanied by disturbances in sleep-wake rhythms as well as alterations in mood, behaviour and daily activities ^1, 2^. In this context, it is well documented that the neurons of the central circadian pacemaker in the brain, the suprachiasmatic nuclei (SCN), undergo a progressive decline with dementia^3^, resulting in a decrease in circadian rhythm amplitude and fluctuations in circadian phase with consequently weaker entrainment to the 24-h day ^4^. These changes in the SCN, together with the natural age-related changes in the visual system, contribute to deterioration in mood, sleep-wake cycles and behaviour, and together, these symptoms increase the need for more intensive care and medication prescription ^5^. Clinically, exposure to bright light at the appropriate time of day improved behavioural symptoms as well as sleep-wake rhythms in older demented patients ^6–14^ and slowed down cognitive deterioration^12^ (for a systematic review see ^15^). Simulated dawn and dusk at the bedside of institutionalised demented patients was found to advance nocturnal sleep onset by one hour ^16^ and improved mood and wellbeing in the morning ^17^.

Another difficulty among institutionalised older patients is accelerating mortality factors, such as recurrent epidemics of influenza ^18^, one of the 10 leading causes for deaths in the USA. The mortality risk can be decreased by winter flu shots ^19^, which are recommended for older patients in long-term care institutions ^20^. However, influenza vaccination is less effective in older than young individuals ^21^, and the immune response is attenuated with age ^19, 22^. IgA and IgG antibody concentrations decrease with age, with a faster decline of the antibody titers ^20^, especially in very old and frail adults. In consequence, older individuals are likely to be insufficiently protected by vaccination ^23, 24^. More recently, during the COVID-19 pandemic, the older population, especially those individuals who live in institutions (e.g. nursing homes) were most vulnerable to infections, ^25–27^ along with a significantly higher mortality risk when positively tested for COVID-19 ^28^.

There is also a (bidirectional) role of sleep ^29, 30^ and the circadian clock on different immune functions ^31–33^, where the SCN is essentially modulating innate and adaptive immune responses (reviewed in ^34, 35^). Aberrant light exposure (such as with shift work or jet-lag) can induce phase shifts in many circadian clock controlled functions, including immune responses ^36, 37^. The consequences of such desynchronisation are dampened circadian amplitudes of SCN cell expression in animals ^38^. In humans, misaligned rhythms appear to be linked to serious health problems such as higher risk for cardiovascular, metabolic and neurodegenerative disease, cancer and impaired immune function ^31^. Thus, indirectly, decreased circadian amplitudes are likely to impair the ability to respond to infections. So far, few studies have addressed the direct impact of ocular light exposure on the human immune system ^39^. One study showed that exposure to continuous bright light during daytime (i.e. polychromatic white electric light; 5000 lx; between 6:30am and 10:30pm) significantly increased unspecific salivary IgA antibody formation in healthy young subjects ^40^. The authors concluded that brighter light exposure during daytime activated a greater immune response in human mucosa cells ^40^.

Enhanced daytime light exposure can increase circadian amplitude of rest activity cycles in older, institutionalised individuals (including those suffering from dementia) ^7, 13^. It is not known whether daily exposure to bright light might also ameliorate adaptive immune responses either indirectly by increasing zeitgeber strength to stabilise the circadian system or directly via an acute action on brain areas regulating innate immune responses. One example of such an adaptive immune reaction is the production of specific antibodies in response to an influenza virus vaccination. An increased specific antibody response to vaccination after enhanced environmental lighting conditions would ameliorate virus protection and thus directly improve general health in these vulnerable patients

Thus, the aim of this study was to assess vaccine responses to three different virus stains of the annual vaccine in a cohort of institutionalised patients with severe dementia. We hypothesised that patients with higher daily light exposure over several weeks would show increased antibody titers in response to the influenza vaccination than patients with lower daily exposure.

## Methods and Participants

### Study design

The cross-sectional study took place during 8 weeks in fall/winter in a Nursing Home in Wetzikon (Zurich, Switzerland) in 2012. Patients in twelve different wards spent time in dayrooms equipped with conventional or ‘dynamic lighting’ systems, where illuminance and correlated colour temperature varied across the day ^11^. We have previously reported the results from patients’ daily activities, agitation, alertness mood, quality of life (from questionnaires assessed by staff members), rest-activity cycles, sleep (derived from activity monitors), and melatonin concentrations (from saliva samples) ^11^. Here we present the results of the specific antibody responses to the annual influenza vaccination.

### Participants

The study group comprised patients over 50 years with one of the following dementia diagnoses (according to DSM-IV): vascular dementia, Alzheimer dementia, frontotemporal dementia, Parkinson’s dementia or mixed forms of dementia. Initially, 104 patients were included in the study ^11^. Here, only patients who had worn the activity watches, received the annual flu shot, and gave two blood samples, were included in the analysis (n = 80; see Supplemental Table S1 for exclusion details). None of the patients was visually blind. Mean age was 78.3 ± 8.9 years (54 women, 26 men; range 55 - 95 years). Ethical approval for all study procedures was obtained from the local Ethical Review Board (KEK, Zurich, Switzerland, protocol # KEK-ZH 2012-0059, now SWISSMEDIC). Written informed consent for study procedures was obtained from family members or legal representatives prior to study begin.

### Individual light exposures

Individual light exposure (illuminance) was recorded via wrist-worn activity monitors equipped with a calibrated light sensor (Motion Watch 8 ®, Camntech, UK). All recordings were downloaded weekly to a PC and visually inspected. Light and rest-activity data was visually inspected and edited by trained assistants (see ^11^ for a detailed description). In brief, if there was a 24-h day with a gap of recorded rest-activity lasting less than 3 hours, this gap was edited with the 24-hour mean of this person. If there was a gap with no rest-activity data for more than 3 hours, that 24-h period was excluded from further analyses. The same criteria were also applied for the light recordings except that if a 24-h period contained rest-activity, but no light data for more than 3 hours (due to coverage of the sensor by sleeves), only the light data was not used, and if the gap was less than 3 hours, the light data was also interpolated with the 24-h mean of that day.

Based on the averaged individual light exposure, two subgroups were created retrospectively, as assessed from wrist worn light sensors (Motion Watch 8, Camntech, UK), similarly to what was done for the previously reported data ^11^). For this purpose, median illuminance across 8 weeks between 8:00 and 18:00 was calculated for each participant which resulted in 426.1 ± 304.4 lux (mean ± SD). In a next step, a median split of these data (n = 80) resulted in a group with higher mean light exposure (= high light group, i.e., > 392.65 lx; n = 40, 27 women, 13 men) and a group with lower mean light exposure (= low light group, i.e., < 392.65 lx; n = 40, 27 women, 13 men,). The two light exposure groups did not differ in age (low light group = 79.0 ± 9.2 years; high light group = 77.7 ± 8.6 years; p = 0.6; Wilcoxon 2-sample test), or cognitive impairment, as assessed by the Severe-Mini Mental State Evaluation (S-MMSE before the flu shot ^41^). The S-MMSE score was 8.1 ± 1.5 for the low light group and 7.8 ± 1.5 for the high light group (p = 0.93 Wilcoxon 2-sample test).

### Rest-activity cycles and sleep

From the wrist worn monitors, circadian rest-activity data were derived as described in ^11^ and above. All data which included 24-h days per patient underwent a non-linear circadian regression analysis ^13, 42^, resulting in the following variables: inter-daily stability (IS), inter-daily variability (IV), and relative amplitude (RA). The RA is defined as the ratio of the 10 hours with highest activity (M10) relative to the 5 hours with lowest activity (L5) per 24 hours. Sleep variables from nighttime sleep episodes were determined by the software Sleep Analysis v7.23 (Camntech UK). Bed- and wake times were assessed as described in ^11^ and sleep variables were re-analysed for the 80 participants: habitual bedtime, wake time, time in bed, sleep duration, sleep efficiency (ratio between sleep duration: time in bed) and fragmentation index

### Blood samples and influenza vaccination

Two blood samples were obtained by standard venous puncture, one immediately before and one approximately 4 weeks after the annual influenza vaccination. A total of 8 ml blood was drawn from each patient by professional staff members of the nursing home before noon. The blood sample was sent to an external laboratory for general analyses (Medica AG, Zürich, Switzerland; Table 1). For the specific antibody titer analysis (Prof. A-C Siegrist, University of Geneva), 4 ml whole blood was coagulated within 1 hour after the sample was taken at room temperature, centrifuged, and the serum pipetted into tubes (Eppendorf ®) and immediately frozen at −20 °C before sending to the University of Geneva for hemagglutination inhibition assays (HIA) ^43^.

**Table 1:** Circadian and sleep variables (derived from activity monitors) for low and high light exposure (low LE, high LE) groups (mean, SD in brackets; n =80). IS = Inter-daily stability; IV = intra-daily variability; L5 = 5 hours with lowest activity; M10 10 hours with highest activity; RA = relative amplitude (see Ref. ^42^ for more details; BT = habitual bedtime (h); WT = habitual waketime (h); TIB = Time in bed (h); Wake = waketime during scheduled sleep (h); Sleep duration (h); SE = sleep efficiency (%); sleep time / TIB × 100); Fragmentation index (dimensionless). * = p < 0.05 between light exposure groups.

| Variable | Low Light<br>Exposure Group | SD | High Light<br>Exposure Group | SD |
| --- | --- | --- | --- | --- |
| **IS \*** | 0.33 | (0.15) | 0.38 | (0.14) |
| **IV** | 1.05 | (0.30) | 1.18 | (0.38) |
| **L5** | 705.77 | (591.84) | 655.15 | (625.49) |
| **M10** | 3969.53 | (3263.18) | 4241.26 | (3756.49) |
| **RA \*** | 0.66 | (0.18) | 0.72 | (0.15) |
| **BT (h)** | 19.50 | (1.00) | 19.57 | (1.31) |
| **WT (h)** | 8.24 | (0.71) | 8.10 | (0.79) |
| **TIB (h)** | 12.71 | (1.25) | 12.52 | (1.73) |
| **Wake (h)** | 2.14 | (1.26) | 1.84 | (1.09) |
| **Sleep duration (h)** | 10.28 | (2.26) | 10.40 | (2.48) |
| **SE (%)** | 80.09 | (12.08) | 82.23 | (10.90) |
| **Fragmentation Index** | 50.81 | (24.87) | 46.69 | (18.86) |

The vaccination was performed in week 44 (i.e. between Oct. 29^th^ and Nov. 4^th^, 2012; according to national recommendations in Switzerland), except for three patients who received the first blood sample and the flu shot in week 46 and the second blood sample at the end of week 50 (due to an acute infection in week 44). The trivalent vaccine Fluarix ® (GlaxoSmithKline Biologicals, UK) was applied via intramuscular injection in the patients’ upper arm by the nursing home staff. The vaccine contained attenuated virus particles against influenza A virus (H3N2, H1N1) as well as against influenza B virus (IB). The strains matched the World Health Organization (WHO) recommendations for the influenza season 2012/2013. Immune responses were calculated only in patients who received the flu shot and had no acute infection (i.e. less than 15’000 Leucocytes/μl) which resulted in 80 patients. Patients who had very high antibody titer concentrations (> 1000) in the pre-vaccination blood sample for one of the three virus strains were also excluded for the analysis of that virus strain, which was the case in two patients for the H3N2 antibody titer, and in one patient for the H1N1 antibody titer.

### Statistical analysis

For rest-activity cycles and sleep, the same analyses as described in ^11^ were performed for the subset of participants undergoing the antibody titer analysis (n=80). For influenza vaccination, the pre- and post-vaccination antibody titers and the ratio (pre-vaccination/post-vaccination) were used to compare the two light exposure groups. The fixed factors LIGHT EXPOSURE GROUP (low light vs. high light), AGE as a categorical factor (on dichotomized variables derived from median split of age, i.e. < 80.0 or >79 y), and SEX as well as their interactions were added to the model. Subject was added as random factor. For general blood variables the repeated factor SESSION (i.e. pre- and post-vaccination blood sample) was included. Statistical comparisons were performed with a generalized linear mixed model (GLIMMX; with a lognormal distribution if the data was not normally distributed) by using the Software Package SAS (SAS Institute Inc., Cary, NC, USA; v 9.4). Degrees of freedom were determined with the Satterthwaite approximation and post-hoc comparisons were performed with the Tukey-Kramer test (corrected for multiple comparisons).

In order to compare the overall immune protection against the influenza virus, we determined geometric mean titers (GMTs) for pre- and post-vaccination values ^43^. The differences between high and low light exposure groups for post-vaccination GMTs were compared by survival analyses (Kaplan-Meier on log-ranked values; Sigma Plot v11.0, Statsoft Software Inc). For both light exposure groups, seroprotection rates (which are defined as percentage of patients with post-vaccination antibody titer ≥ 1:40) and seroconversion rates (described in reference ^43^), defined as percentage of patients with 4-fold increase of pre-vaccination GMT titer, were also calculated.

## Results

### Rest-activity cycles and sleep

Rest-activity cycles revealed a significantly higher inter-daily stability (IS) and relative amplitude (RA) in the high light group compared to the low light group (Table 1; main effect of LIGHT EXPOSURE GROUP; p < 0.05; n = 80; F_1,72_ > 4.2; p < 0.05). In general, IS and RA were higher in women than men (IS: women: 0.39 ± 0.03, men: 0.30 ± 0.02; means, SD; RA: women: 0.72 ± 0.15, men: 0.66 ± 0.18; main effect of SEX; p = 0.004). Activity of the five hours with lowest daily activity (L5) was significantly higher in men (861 ± 572) than in women (594 ± 607; main effect of SEX; F_1,72_ = 8.6; p < 0.05), and men also had earlier wake times than women (wake times men: 7.9 h ± 0.7; women 8.3 h ± 0.8; main effect of SEX; F_1,72_ = 4.0; p < 0.05), which did not result in any other statistical differences of sleep duration or time in bed between men and women. There were no significant main effects or interactions for the remaining circadian or sleep variables.

### Blood variables

There was no difference between the two light exposure groups for any of the blood variables between in the pre- and the post-vaccination samples (p > 0.13, Table 2). In general, the CD4/CD8 ratio, C-reactive protein (CRP), leucocyte count and percentage of neutrophil lymphocytes were higher in the pre- than the post vaccination sample (main effect of SESSION; p < 0.05; Table 2), while the absolute lymphocyte count and percentage were higher in the post- than the pre-vaccination sample (p < 0.05). For the younger subgroup (see statistics) of patients, the erythrocyte counts (EC), haemoglobin (HG), haematocrit (HK), mean corpuscular haemoglobin concentrations (MCHC) and lymphocytes (in % from automated processing) were significantly higher than in the older patient group (main effect of AGE; p < 0.015).

**Table 2:** Blood analyses before and 6 weeks after the influenza vaccination. CRP = C-reactive protein, LC = leucocytes, EC = erythrocytes, HB = haemoglobin, HK = haematocrit, MCV = mean corpuscular volume, MCH = mean corpuscular haemoglobin, MCHC = mean corpuscular haemoglobin concentration, TC = thrombocytes, LUC = large unstained cells, FACS = fluorescence activated cell sorting (CD3, CD4, CD8); n = 80; * = p<0.05; main difference between pre- and post-vaccine session.

| Blood Marker | Pre-vaccination (SEM) |  | Post-vaccination (SEM) |  |
| --- | --- | --- | --- | --- |
| CD4 (count /μl) | 879.23 | (33.17) | 892.61 | (36.09) |
| CD8 (count /μl) | 435.74 | (35.63) | 448.29 | (33.43) |
| CD4/CD8 Ratio * | 2.76 | (0.19) | 2.65 | (0.18) |
| Lymphocytes (count /μl) * | 1775.00 | (86.17) | 1881.01 | (72.40) |
| CRP (mg/l) * | 10.40 | (2.03) | 6.59 | (1.03) |
| LC (count /μl) * | 7110.00 | (243.31) | 6602.50 | (189.08) |
| EC (G/l) | 4.37 | (0.06) | 4.32 | (0.05) |
| HB (g/l) | 130.09 | (1.70) | 128.85 | (1.57) |
| HK (l/l) | 0.40 | (0.005) | 0.39 | (0.004) |
| MCV (fl) | 90.64 | (0.52) | 90.94 | (0.48) |
| MCH (pg) | 29.77 | (0.19) | 29.85 | (0.16) |
| MCHC (g/l) | 328.40 | (0.87) | 328.43 | (0.78) |
| TC (G/l) | 292.91 | (8.60) | 284.41 | (8.24) |
| Neutrophil (%) * | 62.71 | (1.17) | 59.92 | (1.13) |
| Eosinophil (%) | 3.66 | (0.26) | 4.12 | (0.41) |
| Basophil (%) | 0.53 | (0.03) | 0.54 | (0.03) |
| Monocytes (%) | 6.07 | (0.19) | 6.17 | (0.15) |
| Lymphocytes (%) * | 23.94 | (0.94) | 26.50 | (1.00) |
| LUC (%) | 3.08 | (0.27) | 2.76 | (0.17) |
| FACS (count /μl) | 1331.20 | (57.60) | 1358.33 | (55.55) |

### Influenza vaccination response

Pre- and post-vaccination antibody titers did not show significant differences between the light exposure groups (p > 0.07). However, the post/pre-vaccination ratios revealed a significantly higher increase in antibody titer for the H3N2 virus strain in the high than the low light group (Table 3a; F_1,70_ = 6.8; p = 0.01; Figure 1). On visual inspection, the IB post/pre-vaccination ratio seems similar to the H3N2 but the difference did not reach significance. It revealed a trend (p=0.08) for slightly higher antibody titer in the high light exposure group (for significant effects with AGE and SEX see Supplemental Results).

**Table 3a:** Absolute influenza antibody titers pre- and post-vaccination for three virus strains: H3N2 (n=78), H1N1 (n=79) and IB (n=80) as well as ratio (post/pre) for the low and high light group); means and (SEM). * = significant differences between ratio of the low and the high light exposure group (p < 0.05; main effect of light exposure group).

| **Titer** | **Low Light Exposure Group** |  | **High Light Exposure Group** |  |
| --- | --- | --- | --- | --- |
| **Pre (H3N2)** | 140.8 | (12.3) | 150.3 | (25.3) |
| **Post (H3N2)** | 421.3 | (69.2) | 627.5 | (146.3) |
| **Ratio (H3N2) \*** | 3.4 | (0.6) | 9.9 | (3.2) |
| **Pre (H1N1)** | 39.5 | (8.4) | 62.5 | (12.3) |
| **Post (H1N1)** | 477.5 | (93.8) | 401.6 | (112.6) |
| **Ratio (H1N1)** | 31.1 | (7.9) | 26.7 | (9.4) |
| **Pre (IB)** | 65.4 | (9.4) | 56.0 | (9.4) |
| **Post (IB)** | 232.6 | (61.2) | 201.9 | (29.5) |
| **Ratio (IB)** | 5.6 | (1.6) | 9.7 | (2.4) |

**Figure 1.**
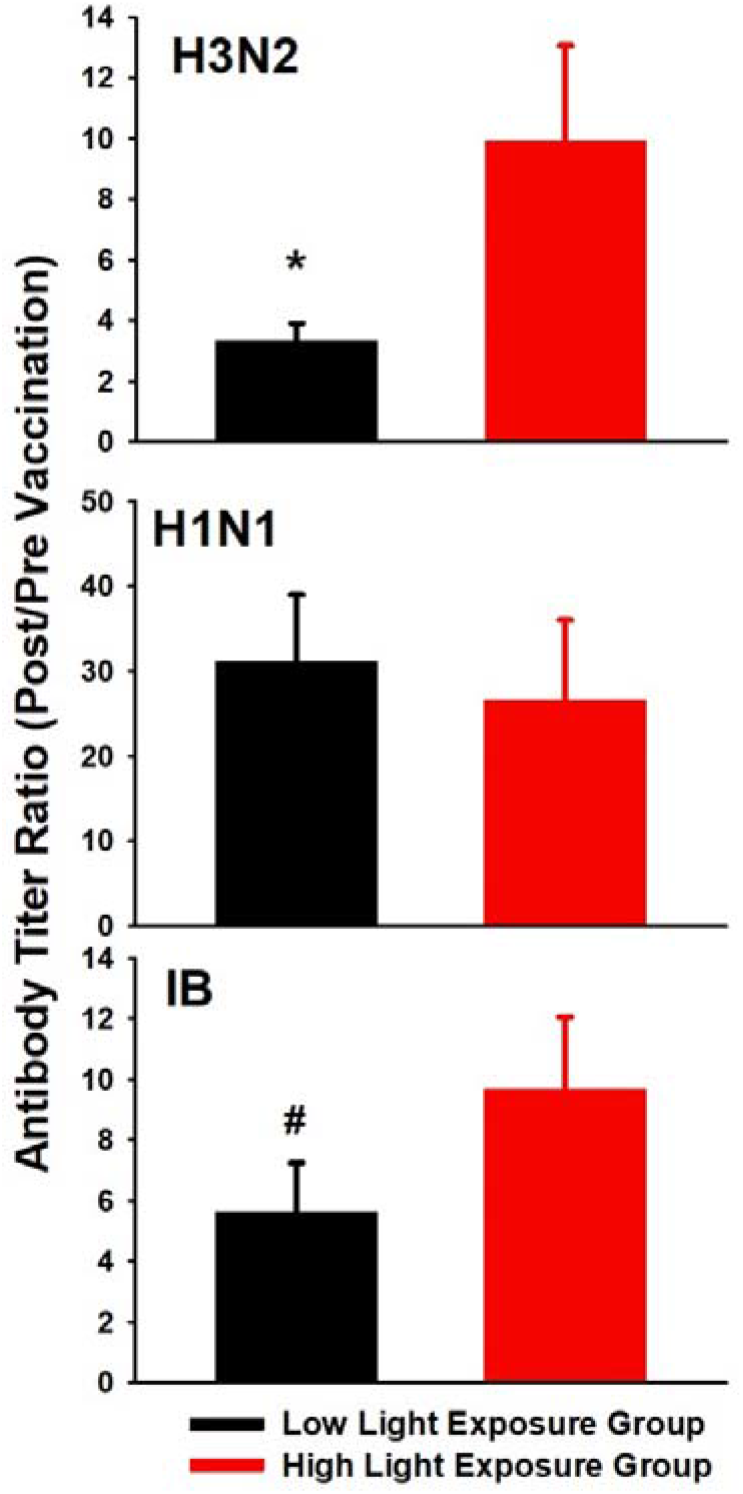
Mean values (+ SEM) for antibody titer ratios (post-vaccination/pre-vaccination) for all three influenza strains and both sub-groups of patients [high (red bars) vs. low light exposure group (black bars)]: H3N2 (n = 78); H1N1 (n = 79); IB (n = 80). *: p = 0.01 (main effect of light exposure group) and trend: #: p = 0.08.

The geometric mean antibody titer (GMT) for pre- and post-vaccination responses are shown in Table 2b for both light exposure groups. From a cohort perspective, the post-vaccination seroconversion rates (i.e., the percentage of patients with a GMT greater or equal 40) were at least 75 % (Supplemental Table S2). The seroprotection rates, which reflect a 4-fold increase of the GMT was greater or equal to 34 % for all three virus strains. When performing a survival analysis on the reverse cumulative distributions of GMTs with the low and high light patient group (Figure 2), there were no significant differences for the three influenza virus strains (Kaplan-Meier on log-ranked values; p > 0.4).

**Figure 2.**
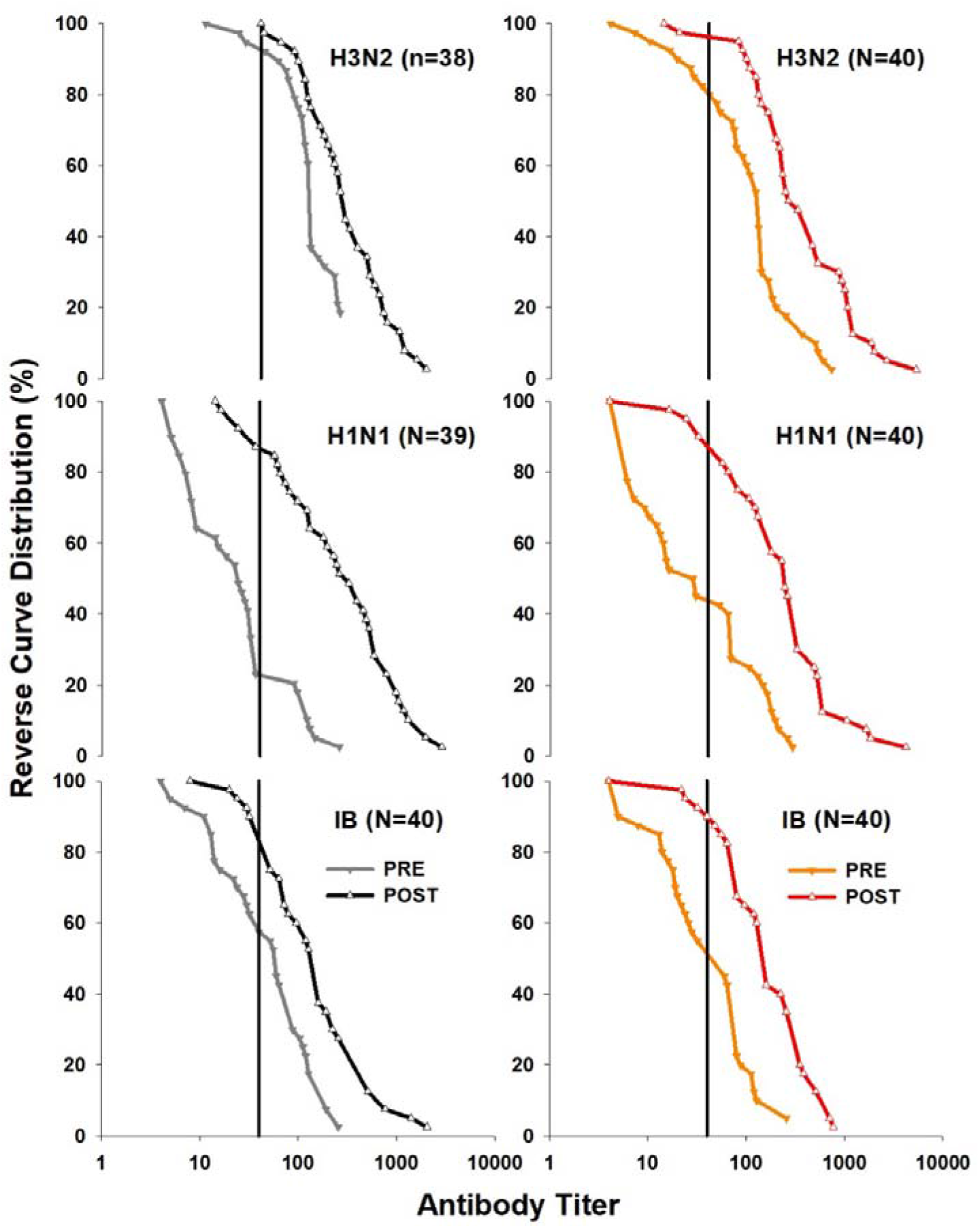
Reverse curve distribution plots from geometric mean antibody titers (GMT) for pre-vaccination (dashed lines) and post-vaccination (solid lines) samples and the three influenza virus strains (H3N2, upper graph; H1N1, middle graph; IB, lower graph). The data is expressed in percentage for both light exposure groups of patients separately (left panel and grey or black symbols = low light exposure group; right panel and orange or red symbols = high light exposure group; filled triangles down, grey or orange symbols = pre-vaccination antibody titers; open triangles up, black or red symbols = post-vaccination antibody titers). The vertical line in each graph represents the threshold for GMT titers of seroprotection by the influenza vaccination (i.e, a GMT ≥ 40).

## Discussion

A group of dementia patients with long-term brighter daytime light exposure showed significantly greater circadian inter-daily stability and higher relative amplitude of circadian rest-activity cycles along with higher antibody production in response to the influenza virus strain H3N2 than the patient group with lower bright light exposure. The seroconversion and seroprotection rates, which are standard criteria to evaluate the effectiveness of a vaccination within a cohort, showed that both light exposure groups were well protected against the influenza virus. However, in highly vulnerable patients, any additional benefit at the individual immunological response level is desirable. Even more so because the effects could have been indirectly conveyed via the improved circadian rest-activity cycles. Indeed, this assumption is corroborated by the significantly higher RA from rest-activity cycles in those participants with enhanced light exposure. A higher relative amplitude with greater inter-daily stability may have repercussions on other health related parameters, as shown for a variety of diseases (reviewed e.g. in ^44, 45^). Although only one influenza strain showed significantly increased response, one other strain (IB) trended to increase in a similar way (p = 0.08).

The results from blood factors measured before and after the influenza vaccination were all in the normal range and revealed some vaccination effects (e.g., lymphocyte count) and some lower values for the older subgroup (e.g., erythrocytes and haemoglobin). There was no statistically significant difference in any blood factor between the light exposure groups, including those involved in T-cell related immune responses (e.g., the CD4 and CD8 cells and the CD4/CD8 ratio). Any differences in antibody responses were rather conveyed through the humoral and adaptive immunity driving cells, the B-lymphocytes.

There is some evidence that more daytime light (= increased Zeitgeber strength) can lead to increased amplitude for example of melatonin secretion profiles in older institutionalised individuals ^46^, or stabilised rest-activity profiles in Alzheimer patients ^13^. Therefore, it is tempting to speculate that the higher immune responses in our patient group with brighter light exposure were also indirectly facilitated by overall higher circadian rhythm stability of central (and peripheral clocks).

There are other influencing factors which may also indirectly contributed to our results, e.g., it is well known that stress, emotional or other biopsychological and social aspects (bidirectionally) affect acute and long-term immune responses ^47, 48^. In our sample, we previously reported more expressions of positive emotions, greater alertness and higher quality of life over an 8 week observation period in the group with higher light exposures ^11^. Therefore, since these positive findings occurred in the same patient group, it may be that brighter light exposure improved immune responses not only by increasing circadian stability but also by positive emotions such as improved mood, greater alertness and quality of life ^11^.

Beneficial non-visual effects of brighter light exposure can obviously be best provided by natural daylight (with appropriate UV-protection of skin and eyes), but also by improved electrical lighting systems^7, 8, 10, 12, 49, 50^. It is well known that light through the skin stimulates innate and adaptive immune responses, thus preventing different diseases through Vitamin D production from Ultraviolet B (UVB) solar radiation ^51, 52^. In our dementia patient cohort, we used wrist-worn light sensors which continuously measured illuminance. A limitation of our study may be that from these measures we could not disentangle whether participants were outside and exposed to direct sun (and UVB) or spending time inside, where window glazing absorbs UVB radiation.

In summary, there may be underestimated benefits from regular brighter and natural light exposures in older and frail individuals: better circadian entrainment, mood, rest-activity and immune functions. Our findings provide some evidence that in patients with dementia, long-term brighter light exposure may foster increased antibody titer production in response to the annual influenza vaccination, either directly or via circadian rhythm stabilisation. Further demonstration of positive effects of light to boost immune functions would open a whole new area of research with wide applications. There is a need to determine the optimal timing, duration and qualities of light required.

In summary, we have some first evidence that brighter daily illuminance levels, provided by natural daylight or improved electrical lighting, might be beneficial for demented patients in modulating specific immune responses. To disentangle possible causal relationships, prospective clinical studies investigating light exposure effects on these variables are warranted.

## Acknowledgments

We thank all care givers and professionals of the Nursing home ‘Das Heim’, especially Michael Schmieder and Katharina Bieler, for without their professional help and support, the study would not have been possible. We thank Anne-Claire Sigrist and team of for great help with the antibody titer assays and Adriano Fontano for his initial advice with the immunological variables. We are also grateful to Markus Oetiker from Zumtobel Licht AG (Dornbirn, Austria) for help with programming the dynamic light settings. We also thank Timo Fuhrman with the data collection, light measurements and him and Daniel Hulliger for his valuable help with data processing. We also acknowledge Licht Zumtobel AG (Austria), Medica AG (Switzerland) and Camntech (UK) and Salimetrix (UK) for giving us a discount price on their products.

