## Supplemental_File for "Enhancing daily light exposure increases the antibody response to influenza vaccination in patients with dementia"

### Supplemental Results

#### Influenza vaccination responses: effects of AGE, SEX

Pre- and post-vaccination antibody titers did not show significant differences between the light exposure groups ($p > 0.07$). The interactions of LIGHT EXPOSURE GROUP with SEX and/or AGE were not significant. The H1N1 antibody titer of the pre-vaccination was significantly higher in the older (73.2 ± 12.4; n = 41) than the younger subgroup [(27.4 ± 6.2; n = 38) main effect of AGE; $F_{1,71} = 8.9$; $p = 0.004$)]. For the IB antibody titer, there were significant interactions between AGE and SEX such that pre-vaccination antibody titers were lower for older men than women (older men 35.8 ± 13.2, n = 11; older women 81.3 ± 10.4, n = 31; $F_{1,72} = 6.5$; $p = 0.01$). IB antibody titer ratios were significantly higher for older men only in the high light exposure group (26.3 ± 10.6; n = 5) than women of the same group (mean 6.2 ± 2.9 SD; n = 17; interaction LIGHT EXPOSURE GROUP x SEX x AGE; 5.6 ± 1.8; $F_{1,72} = 9.7$; $p = 0.003$).

**Supplemental Table S1:**

| **Participant excluded (#)** | **Reason** |
| --- | --- |
| 49 | Retraction of consent |
| 66, 76 | Deceased |
| 81 | Transferred to psychiatry (> 3 weeks) during an acute episode |
| 73, 82, 99 | Eye disease (Glaucoma, Retinitis Pigmentosa) |
| 8, 40, 70 | No activity watch data |
| 6, 29, 38, 52, 84 | Insufficient light data |
| 3, (8) 21, 39, (40), (52), 54, 55, 59, 91 | Not 2 blood samples and/or no influence vaccination (in brackets = patients already excluded from analysis) |
| 7, 44, | Leucocyte cell count > 15’000 / µl (likely due to acute infection) |

**Supplemental Table S2:**

| Strain | Groups | GMT | Seroprotection Rate | Seroconversion Rate |
|---|---|---|---|---|
| **H3N2 (n=78)** | **Pre: low** | 116.51 | 92 | - |
| | **high** | 88.82 | 80 | - |
| | **Post: low** | 272.15 | 100 | 34 |
| | **high** | 309.88 | 95 | 38 |
| **H1N1 (n=79)** | **Pre: low** | 19.97 | 21 | - |
| | **high** | 25.11 | 43 | - |
| | **Post: low** | 218.35 | 85 | 74 |
| | **high** | 171.51 | 83 | 73 |
| **IB (n=80)** | **Pre: low** | 40.10 | 58 | - |
| | **high** | 32.67 | 45 | - |
| | **Post: low** | 112.26 | 75 | 38 |
| | **high** | 128.41 | 90 | 43 |

**Supplemental Table S2:** Shows the geometric mean antibody titers (GMT) for each influenza strain before and after vaccination and for both sub-patient groups (low light, high light group respectively). The sero-protection rate indicates the portion of the cohort with a post-vaccination titer ≥ or 40 (in %). The seroconversion rate shows the proportion of the cohort with 4-fold GMTs (in %).
